# Efficient analysis of large datasets and sex bias with ADMIXTURE

**DOI:** 10.1101/039347

**Authors:** Suyash S. Shringarpure, Carlos D. Bustamante, Kenneth Lange, David H. Alexander

**Affiliations:** Department of Genetics, Stanford University, Stanford, California, USA; Department of Biomathematics, UCLA, Los Angeles, California, USA; Pacific Biosystems, Palo Alto, California, USA

**Keywords:** supervised learning, reference panels, pedigrees, sex-chromosome, sex bias, ancestry ineference, admixture

## Abstract

**Background**: A number of large genomic datasets are being generated for studies of human ancestry and diseases. The ADMIXTURE program is commonly used to infer individual ancestry from genomic data.

**Results:** We describe two improvements to the ADMIXTURE software. The first enables ADMIXTURE to infer ancestry for a new set of individuals using cluster allele frequencies from a reference set of individuals. Using data from the 1000 Genomes Project, we show that this allows ADMIXTURE to infer ancestry for 10,920 individuals in a few hours (a 5x speedup). This mode also allows ADMIXTURE to correctly estimate individual ancestry and allele frequencies from a set of related individuals. The second modification allows ADMIXTURE to correctly handle X-chromosome (and other haploid) data from both males and females. We demonstrate increased power to detect sex-biased admixture in African-American individuals from the 1000 Genomes project using this extension.

**Conclusions:** These modifications make ADMIXTURE more efficient and versatile, allowing users to extract more information from large genomic datasets.

## Todo list

### Background

The ADMIXTURE program [1] estimates individual ancestry proportions for admixed individuals from genomic datasets. It uses a likelihood model [2] that assumes the genotype *n_i_j* for individual i at biallelic SNP *j*, which represents the number of type “1” alleles observed, is generated by binomial sampling from a weighted sum of ancestral allele frequencies. For each individual, the weights are given by the proportions of ancestry derived from each ancestral population. Given *K* ancestral populations, genotypes are sampled as 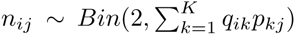 where *q_ik_* the fraction of individual *i*’s ancestry attributable to population k and *p_kj_* is the frequency of the type 1 allele at SNP *j* in population *k*. ADMIXTURE maximizes the resulting biconcave log-likelihood (Equation 1) using a block relaxation algorithm.

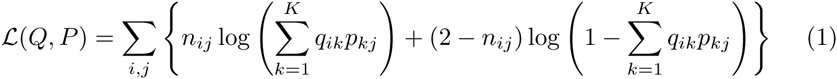

We describe two extensions to the ADMIXTURE program that accelerate the analysis of large datasets and enable ancestry estimation for sex chromosomes. The first extension (“projection”) allows ADMIXTURE to estimate ancestry for a new set of individuals using ancestral populations from an earlier ADMIXTURE run. It enables efficient inference of ancestry on large genomic datasets using ancestral populations estimated from reference panels like the 1000 Genomes Project. It can also be used to correctly infer individual ancestry in pedigrees. The second extension allows ADMIXTURE to model the log-likelihood for haploid chromosomes. This can be used to correctly estimate ancestry on sex chromosomes and therefore estimate sex bias in ancestry between the autosomes and sex chromosomes. We demonstrate the utility of these extensions using data from the 1000 Genomes Project [3] and the HapMap Project [4].

## Implementation

### Projecting new samples on existing population structure

A number of large genome-wide datasets of human populations such as the HapMap Project, 1000 Genomes Project etc. are now publicly available. Many studies (e.g. [5]) use these datasets as reference panels in combination with the study sample to estimate individual ancestry using ADMIXTURE since these large datasets summarize worldwide human population structure. For study samples which do not include a novel population, an efficient way of estimating individual ancestry is to “project” the new samples on to the population structure learned from the reference panels. This is intuitively similar to the projection operation used in principal components analysis, though the mathematical details differ. We extended the ADMIXTURE code to allow loading of trained models (the .P files with cluster allele frequencies). For two datasets with the same set of SNPs, clusters can be learned using the unsupervised mode of ADMIXTURE on the first dataset and ancestry proportions can be inferred for the second dataset using these learned clusters. The same approach can be used to infer ancestry on a set of related individuals. First, we infer the largest set of unrelated individuals in the dataset using pedigree information or methods such as PLINK [6], KING [7] or PRIMUS [8]. Then, ADMIXTURE is run on this set in unsupervised mode and the remaining individuals are projected on the resulting population structure.

Mathematically, this requires solving the likelihood maximization problem of Equation 1 with respect to *Q* for a fixed *P*. This a convex problem and can be solved efficiently using the optimization described by Alexander et al. [1].

### Analyzing haploid sex-chromosomes

Admixture between populations is often sex-biased, i.e., different proportions of males and females from the source populations contribute to the admixed populations. In human populations, sex-biased admixture has been observed in African-Americans and Latinos, often using evidence from Y-chromosome or mitochondrial DNA [9, 10, 11]. An alternative way to study sex-biased admixture is to examine individual ancestry estimates on the autosomes vs the sex chromsomes [5, 12]. Therefore, we are interested in inferring individual ancestry using ADMIXTURE on the sex chromosomes, in particular on the haploid X-chromosome in males.

For a haploid sex-chromosome SNP, we assume that hemizygous genotypes are coded as homozygotes for the observed allele. Then, the log-likelihood for a haploid sex-chromosome SNP in an individual is half of that for a homozygous autosomal diploid SNP in Equation 1. We account for this in ADMIXTURE by keeping track of the sex of each individual and the chromosome each SNP belongs to and adjusting the log-likelihood accordingly.

To enable correct handling of haploid sex-chromosomes in multiple species, we implemented the --haploid option, which takes a single colon-separated argument describing the haploid sexes and the haploid chromosomes. For instance, for human data, sex-chromsomes can be supplied as an argument for ADMIXTURE as --haploid=“male:23,24” with 23 and 24 representing the X and Y chromosomes respectively.

## Results

We demonstrate the utility of the newly implemented options using experiments on human genomic datasets.

### Using reference panels for inferring ancestry proportions with projection

We duplicated data from Phase 1 of the 1000 Genomes Project to create a dataset with 10,920 individuals. The data was filtered to include only SNPs with minor allele frequency (MAF) ≥ 5% and thinned for linkage disequilibrium (LD) to have pairwise *r*^2^ ≤ 0.1 in 50 kb windows. We compared the running time and accuracy of two analyses, with the number of clusters (*K*) ranging from 2 to 10:

- **Unsupervised**: Unsupervised ADMIXTURE was run on the entire dataset of 10,920 individuals.
- **Projection**: Unsupervised ADMIXTURE was first run on the original 1,092 individuals from the 1000 Genomes Project and the remaining 9,828 individuals were projected on to the learned population structure.

Each analysis was performed with 5 random starts, with running time limited to 72 hours. All experiments were run on a single core of a server with Xeon E5-2660 processors, using 3.7 GB memory.

Figure 1 shows the comparison of running times for ADMIXTURE on the 10,920 individuals using the two approaches. The projection approach is much faster than unsupervised ADMIXTURE, with speed gains increasing with K, the number of clusters. We find that the ancestry proportions inferred using both approaches are identical.

**Figure 1.**
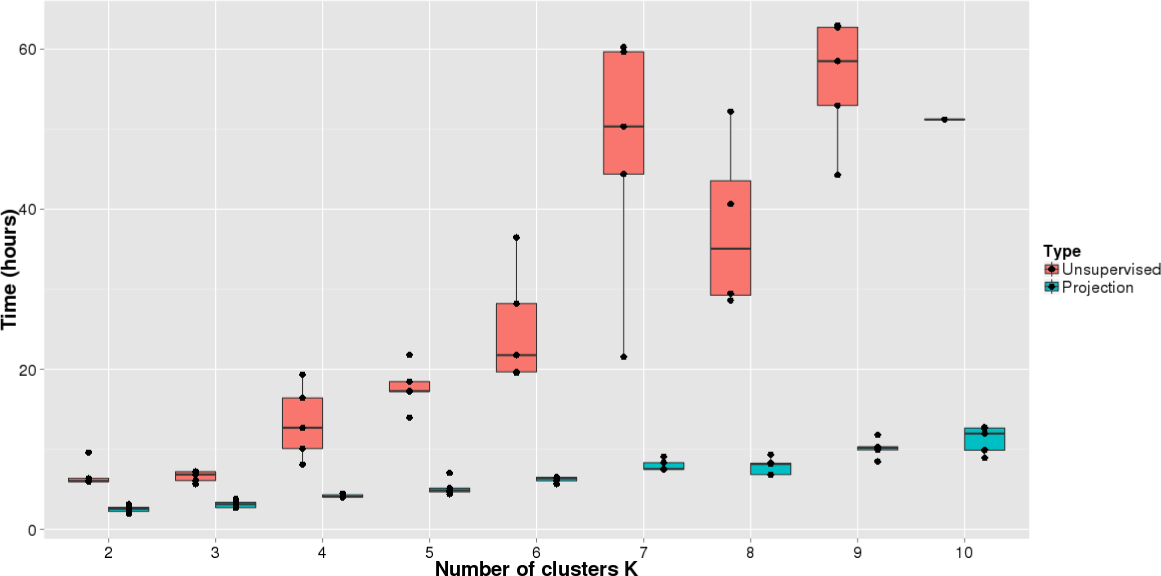
Running time comparison. Running times for ADMIXTURE on a dataset of 10,920 individuals constructed from the 1000 Genomes project.

#### Comparison with iAdmix

The projection step we describe has been recently independently implemented by Bansal et al. [13] in the software iAdmix, using a different optimization algorithm. We compared our ADMIXTURE projection implementation to the iAdmix projection implementation by running unsupervised ADMIXTURE on the first 1,092 individuals from the previous analysis and using the learned allele frequencies to infer ancestry for the remaining 9,828 (copied) individuals by projection using either ADMIXTURE or iAdmix. Figure 2 shows that projection using ADMIXTURE is approximately 4 times faster than using iAdmix^[1]^.

**Figure 2.**
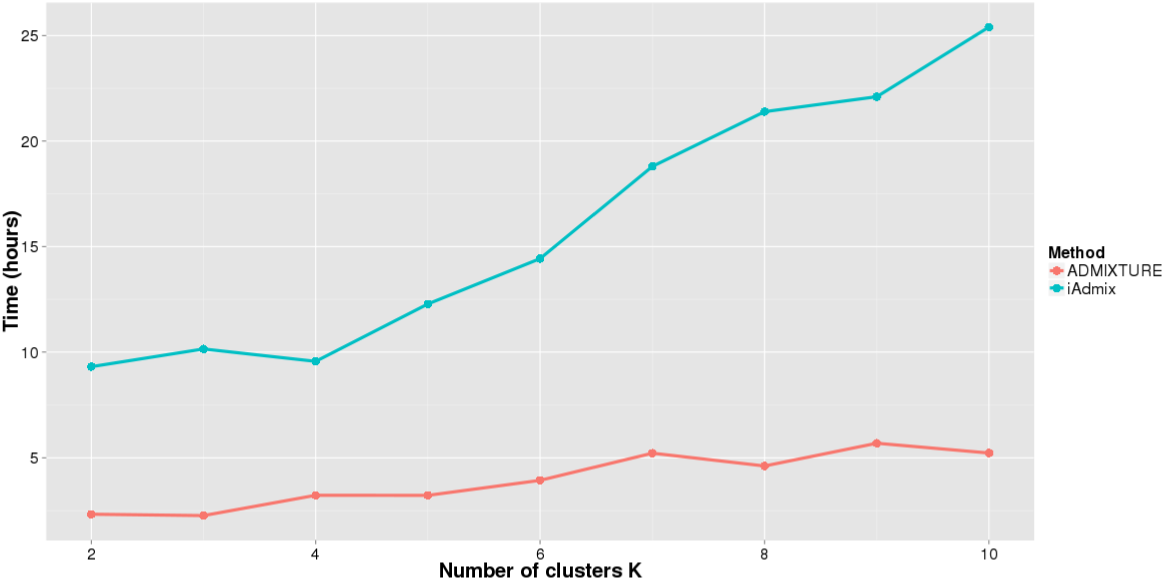
Running time comparison with iAdmix. Running times for the projection step using ADMIXTURE and iAdmix on a dataset of 10,920 individuals constructed from the 1000 Genomes project. Allele frequencies were inferred from the first 1,092 individuals using ADMIXTURE.

### Ancestry estimation for related individuals using projection

ADMIXTURE infers individual ancestry proportion and ancestral population allele frequencies simultaneously in an alternating optimization [1]. Inferring allele frequencies (AF) from related individuals without suitable correction for relatedness can lead to high variance in estimates [14]. We demonstrate that relatedness can affect the inferred population clusters when ADMIXTURE is run on related individuals using the CEPH (Utah residents with ancestry from northern and western Europe, CEU) and Yoruba in Ibadan, Nigeria (YRI) individuals from HapMap Phase 3. We also show how projection can be used to obtain more accurate AF estimates.

We used 165 CEU individuals (112 unrelated and 53 related) and 113 unrelated YRI indviduals to construct a dataset with 278 individuals. After filtering for LD (*r*^2^ < 0.2) and MAF > 0.05, the dataset had 180,591 SNPs. The dataset then was then analyzed using ADMIXTURE with *K* = 2 population clusters in two ways:

- **All individuals**: ADMIXTURE was run on the entire dataset.
- **Unrelated individuals**: The dataset was divided into two sets - one containing only the 225 unrelated CEU and YRI individuals and another containing the 53 related CEU individuals. ADMIXTURE was run on the unrelated set. The related individuals were then projected on the allele frequencies inferred from the unrelated set.

For both analyses, we then compared the inferred allele freqencies for the European components to AF estimates from the Exome Aggregation Consortium (ExAC [15]) data at a common set of 939 SNPs (with frequency between 5% and 95% in ExaAC). We find that European component AF estimates are closer to ExAC allele frequencies for the unrelated analysis (root mean square error=0.040) than for the analysis using all individuals (root mean square error=0.041), with p=0.005 for a one-tailed paired t-test when the squared errors are compared for each SNP. However, this error includes (1) the variance of the estimate due to the sample size from which the AF is estimated and (2) the variance of the estimate due to the relatedness of the samples. Assuming the Exac AF *f* to be the true underlying frequency, a normal approximation for the sample AF *f_n_* estimated from *n* unrelated diploid samples is given by 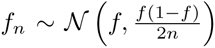.[16] Therefore, we can construct a z-score that accounts for sampling variance as 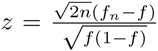. Comparing z-scores, we find that the z-score for the analysis using only unrelated individuals (mean |*z*|=-0.19) is smaller than the z-score for the analysis using all individuals (mean |*z*|=-0.25), with *p* < 2.2e-16 for a one-tailed paired t-test. The z-score using only unrelated individuals also has a smaller variance (var(z)=1.80) than that for the z-score using all individuals (var(z)=2.74). This suggests that the allele frequency estimates from the analysis using unrelated individuals are more accurate than those using all individuals. An alternative way of evaluating the accuracy of estimated allele frequencies is discussed in Supplementary Text Section S1.

### Inference of sex bias from autosomal and X-chromosome ancestry

To demonstrate the utility of ancestry inference on haploid sex chromosomes, we examine sex-biased admixture in the African-American population in the southwestern United States (ASW). We used 1092 individuals from Phase 1 of the 1000 Genomes project including the ASW with populations from Europe, Africa, Asia and the Americas. SNPs were filtered to include only those with MAF ≥ 5% and then thinned for LD to have pairwise *r*^2^ ≤ 0. 1 in 50 kb windows.

Sex bias was analyzed by running ADMIXTURE on the 1092 individuals with *K* = 3 clusters on the autosomes and X-chromosome separately and comparing ancestry proportions for each individual on the two chromosome subsets. If there was no sex-bias during admixture, then the ancestry proportions on the two chromosome sets should be (nearly) equal.

We compared two ways of analyzing sex bias:

- **Females only**: Since ADMIXTURE (without the new --haploid option) requires diploid data, we subset the dataset to 522 females and ran ADMIXTURE on the autosomes and X-chromosome separately.
- **Males and Females**: Using the --haploid option (the X chromosome was denoted haploid in males with --haploid=“male:23”), we ran ADMIXTURE separately on the autosomes and X-chromosome on the entire set of 1092 individuals.

Table 1 shows the results of the analysis. From both analyses, we can see that autosomes have an excess of European ancestry and X-chromsomes have an excess of African and Native American ancestry. To evaluate the significance of the results for each ancestry component (European/African/Native American), we used a paired difference test to compare the means of the X-chromosome and autosomal ancestry proportions. The test statistic is the mean difference in European (for example) ancestry proportion for the X chromosome and the European ancestry proportion for the autosomes for an individual. We estimated p-values using a permutation test with 100,000 permutations (see Supplementary Text Section S2 for details of the permutation procedure). We see that the analysis using both males and females can reject the null hypothesis of identical means (no sex bias) at the 0.05 significance level, while the females-only analysis fails to reject the null hypothesis. From previous work, there is evidence for sex-biased admixture in African-Americans [9, 12, 17]. Thus, including male samples in the analysis of X-chromosome ancestry with the --haploid option improves power to detect sex bias in admixture.

**Table 1.**
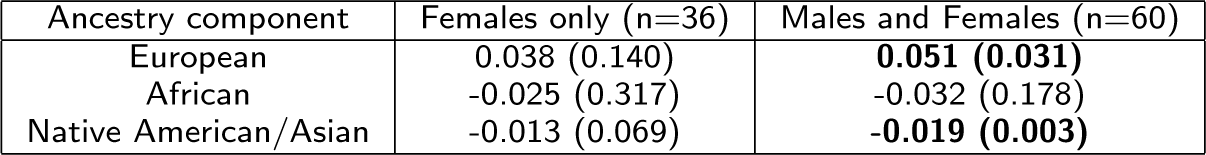
Comparing ancestry proportions for African-Americans on the autosomes and the X-chromosome: Differences in individual autosomal and X-chromosome ancestry proportions are represented by the mean of the difference over all individuals. In parentheses are the raw p-values calculated using 100,000 permutations for a paired difference test comparing the autosomal and X-chromosome ancestry proportions. P-values < 0.05 are shown in bold.

## Discussion

We have described two extensions to the ADMIXTURE program. The projection extension allows ADMIXTURE to estimate ancestry for a new set of individuals using pre-defined ancestral population frequencies (usually from an earlier ADMIXTURE run). This functionality is similar to that implemented in iAdmix [13], which uses a different optimization method, and that implemented by Sikora et al. [18] for ancestry inference for ancient individuals using an expectation-maximization algorithm. This extension enables efficient inference of ancestry on large genomic datasets using ancestral populations estimated from reference panels like the 1000 Genomes Project. The allele frequencies inferred by ADMIXTURE have been used previously to simulate individual genotypes [19, 20]. The resulting individual genomes have been used in subsequent ADMIXTURE [19] or other [20] analyses to enable a “supervised” analysis [21]. Our extension provides an efficient and principled framework for this approach.

The projection approach is useful when a new dataset is strongly unbalanced in its distribution of populations, since an unbalanced dataset can affect the accuracy of ancestry inference [22]. Another advantage of the projection approach is that individual ancestry can be inferred in parallel for each individual. Thus, if a user has access to multiple computers (or a computing cluster), then ancestry can be estimated for hundreds of thousands of individuals in a few hours. Our results on a dataset of 10,920 individuals constructed using the 1000 Genomes project show how projection improves the efficiency of ADMIXTURE. The projection approach can also be used to infer the ancestry of ancient DNA samples, as in Sikora et al [18] and other work. A limitation of the projection approach is that if the projected data contains a novel population which was not present in the initial (training) set, the projection results may not be identical to those obtained from running ADMIXTURE on the combined dataset.

Through experiments on HapMap CEU and YRI individuals, we showed that the projection approach is also useful for accurate ancestry inference on related individuals. This approach allows us to infer allele frequencies for ancestral populations with reduced error. A limitation of this approach is that if the number of founders in a pedigree is small, then the error in allele frequencies estimated from running ADMIXTURE only on the unrelated individuals may be large due to a larger sampling variance. In such cases, the method may not produce more accurate estimates than those obtained by running ADMIXTURE on the entire dataset.

The second extension we have developed correctly models the log-likelihood for haploid chromosomes. This can be used to estimate ancestry on sex chromosomes and thus estimate sex bias in ancestry. Our analysis of sex bias in the ASW African-American population shows that accurate ancestry inference on the haploid X-chromosome in males can improve power of tests for sex bias that use ancestry proportions as a test statistic. While the test we described based on a difference in mean ancestry has a number of limitations (correlated tests, no correction for multiple testing, etc.), it is only intended to demonstrate the advantage of ancestry inference on haploid chromosomes for more power in tests for sex bias and is applicable to other tests of sex bias.

## Conclusions

ADMIXTURE is widely used for analysis of ancestry in genomic datasets. The extensions we have described increase the efficiency of ADMIXTURE and increase its versatility. The projection operation allows more efficient analysis of large datasets by using available reference panels. It also allows analysis of ancestry in pedigrees. Ancestry analysis of haploid sex-chromosomes improves power to detect sex bias in populations using autosomal and X-chromosome ancestry. We expect that with the growing number of populations being sequenced and large amounts of individual-level genotype data being generated, these extensions will make ADMIXTURE more useful to researchers.

### Availability and requirements

Lists the following:

Project name: ADMIXTURE

Project home page: http://www.genetics.ucla.edu/software/admixture

Operating system(s): Linux, Mac OS X

Programming language: C++

Other requirements: None

License: Binaries freely available; source code proprietary

Any restrictions to use by non-academics: None

### Author’s contributions

SSS and KL devised the mathematical details. SSS and DHA implemented the software. SSS and CDB designed the experiments. SSS analyzed data. SSS, KL, CDB and DHA composed the manuscript. The authors have approved the manuscript.

## Acknowledgements

The authors would like to thank Shaila Musharroff for comments on the manuscript and the Stanford Genetics Bioinformatics Service Center for computing resources.

## Supplementary Text for Efficient analysis of large datasets and sex bias with ADMIXTURE

### S1 Evaluating the accuracy of estimated allele frequencies

Another way of measuring the discrepancy between the estimated allele frequencies and the ExAC allele frequencies that takes into account the effect of frequency is to use the binomial deviance, defined for *n* SNPs as *D* = 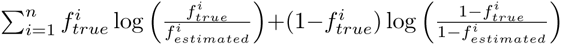 where 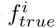 and 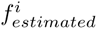 are the true (ExAC) and estimated allele frequencies for the *i^th^* SNP. We find that the binomial deviance for the allele frequency estimates using the unrelated individuals only (7.22) is less than the binomial deviance for the allele frequency estimates using all individuals (7.60), in agreement with our hypothesis that allele frequency estimates from the analysis using unrelated individuals are more accurate than those using all individuals.

### S2 Details of permutation procedure for detecting sex bias in admixture

The test statistic for detecting sex bias in admixture is the mean difference in European (for example) ancestry proportion for the X chromosome and the European ancestry proportion for the autosomes for an individual. Due to the heteroscedasticity of the data, the test statistic does not have a t-distribution. Autosomal ancestry proportion estimates have lower variance than X-chromsome ancestry proportion estimates since they are estimated from a larger number of SNPs. Within the X-chromosome ancestry proportion estimates, estimates for females (with diploid genotypes) have lower variance than estimates for males (with haploid genotypes at the same set of SNPs).

We therefore estimated p-values using a permutation test with 100,000 permutations. For the null distribution of the test statistic, the X and autosome labels were permuted for the ancestries for a single individual. This is equivalent to randomly flipping the sign of the difference in ancestry proportion on the X and autosome for each individual and then recomputing the mean difference.

## Footnotes

[1] We only show results for one replicate since iAdmix produces 130GB of output files for one replicate of such a large dataset.

